# Projecting clumped transcriptomes onto single cell atlases to achieve single cell resolution

**DOI:** 10.1101/2022.04.26.489628

**Authors:** Nelson Johansen, Gerald Quon

## Abstract

Multi-modal single cell RNA assays capture RNA content as well as other data modalities, such as spatial cell position or the electrophysiological properties of cells. Compared to dedicated scRNA-seq assays however, they may unintentionally capture RNA from multiple adjacent cells, exhibit lower RNA sequencing depth compared to scRNA-seq, or lack genome-wide RNA measurements. We present scProjection, a method for mapping individual multi-modal RNA measurements to deeply sequenced scRNA-seq atlases to extract cell type-specific, single cell gene expression profiles. We demonstrate several use cases of scProjection, including the identification of spatial motifs from spatial transcriptome assays, distinguishing RNA contributions from neighboring cells in both spatial and multi-modal single cell assays, and imputing expression measurements of un-measured genes from gene markers. scProjection therefore combines the advantages of both multi-modal and scRNA-seq assays to yield precise multi-modal measurements of single cells.

## INTRODUCTION

In recent years, there has been a surge in the number and size of atlasing efforts across tissues, conditions, and species^1–4^, driven by the high throughput nature of single cell- and nucleus-RNA sequencing (sc/snRNA-seq) technologies. These technologies are now routinely used to generate atlases on the scale of up to millions of cells^3,5–7^, in order to maximize the discovery of novel cell types and characterize the transcriptional heterogeneity of individual cell types within samples. One of the limitations of the sc/snRNA-seq technologies, however, is that they only capture the RNA content of each cell. To address this limitation, there are a growing number of single cell *resolution* assays that simultaneously measure RNA content as well as other cellular annotations and modalities. For example, spatial transcriptomic sequencing assays such as Slide-seq^8^ and LCM-seq^9^ record both the spatial position and RNA measurements from individual spots on a sample. There are also multi-modal assays such as Patch-seq^10^ that measure cellular phenotypes in addition to local RNA content, enabling the identification of connections between molecular and cellular phenotypes of neurons.

However, single cell resolution assays have a major drawback: they often trade off some precision in their RNA measurements in exchange for collecting additional data modalities. In the case of some spatial transcriptome sequencing assays such as LCM-seq or Slide-seq, RNA is extracted from spots of pre-defined size and location on a tissue, leading to individual spots often capturing RNA from multiple cells. Analogously, for Patch-seq, a micropipette is used to puncture brain slices and remove RNA from a target neuron, but RNA from neighboring neuronal or glial cells can be captured as well^11^. For technologies such as MERFISH^12^, in practice only a few hundred genes in the genome can be measured in a tissue. This lack of true single cell, genome-wide RNA measurements can hinder downstream analysis of spatial gene expression patterns or inferring connections between molecular and cellular phenotypes.

Here we present scProjection, a method for projecting single cell resolution RNA measurements onto deeply sequenced single cell atlases, in order to achieve single cell precision from the original RNA measurements. First, we demonstrate our cell type-specific projections capture RNA contributions of component cell types, and importantly that the gene co-expression network of the projected data is consistent with the gene co-expression network of scRNA-seq data from the same cell population. We then illustrate three use cases of scProjection. First, we show scProjection analysis of spatial transcriptomes yields substantially increased detection of cell type-specific spatial gene expression patterns across diverse tissues such as the primary motor cortex and hypothalamic regions of the brain as well as the intestinal villus. Second, we demonstrate scProjection can impute spatial genome-wide gene expression measurements when targeted sequencing of limited numbers of genes via MERFISH^13^ is performed. Finally, we show scProjection can separate RNA contributions from multiple cell types when analyzing Patch-seq data, where RNA measurements are composed of RNA from the target neuron as well as neighboring glial cells. The separation of RNA contributions leads to more accurate prediction of one data modality (electrophysiological response) from another (RNA expression levels). We conclude that integrating deep single cell atlases with single and multimodal cell resolution assays can therefore combine the advantages of both sequencing approaches to study single cells.

## RESULTS

The scProjection model and workflow is illustrated in **Figure 1**. scProjection assumes that one or more RNA samples *x*_*i*_ from a single cell resolution assay are available as input (**Fig. 1a**), as well as a deeply sequenced single cell atlas that profiles the same cell types as the single cell resolution assay (**Fig. 1b**). Typical single cell resolution assays of interest include spatial transcriptome assays such as LCM-seq, Slide-seq or MERFISH, multimodal assays such as Patch-seq, or classical bulk RNA-seq. As output, scProjection simultaneously projects each RNA sample *x*_*i*_ onto each component cell population *k* within the single cell atlas to find the average cell state (expression profile) of that cell type in the sample (*y*_*i,k*_) (**Fig. 1c**), as well as the relative abundance of that cell type (*α*_*i,k*_) (**Fig. 1d**). scProjection therefore balances selecting sets of cell states *y*_*i,k*_ that help minimize reconstruction error of the original RNA measurement *x*_*i*_, with the task of selecting cell states that are frequently occurring in the single cell atlas (e.g. the prior).

**Figure 1.**
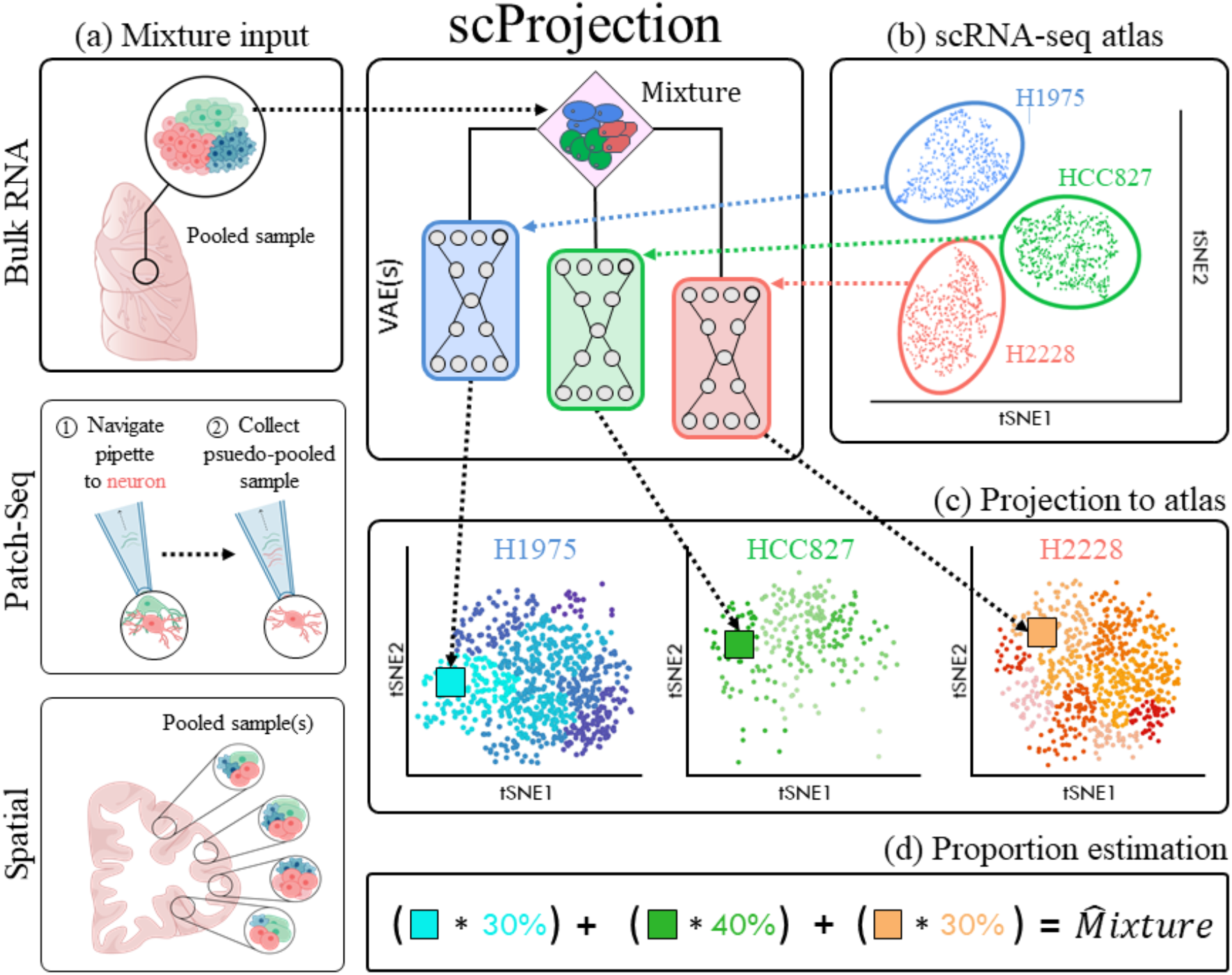
Schematic of cell type projection and abundance estimation with scProjection. **(a)** The primary input to scProjection consists of one or more RNA measurements originating from mixtures of cells assayed using bulk RNA-seq, multi-modal assays or spatial transcriptomics. (**b**) The secondary input to scProjection is a single cell atlas from the same region or tissue as the mixture samples, and is assumed to contain all the cell types present in the mixture samples. For each of the annotated cell types in the single cell atlas, a variational autoencoder is trained to model within-cell type variation in expression. (**c**,**d**) The average cell state for each cell type in a single RNA mixture, along with the relative abundances of each cell type, are estimated by balancing two objectives: (c) selection of an average cell state per cell type that is likely given the single cell measurements for each cell type (the prior), and (d) the joint selection of cell states for each cell type, and abundances, that will lead to the best reconstruction of the original mixed RNA measurements (data likelihood).

scProjection uses individual variational autoencoders^14^ (VAEs) trained on each cell population within the single cell atlas to model within-cell type expression variation and delineate the landscape of valid cell states^15^, as well as their relative occurrence. Here, a valid cell state for a cell type *k* is defined as a genome-wide gene expression profile that has either been directly measured in the single cell atlas, or is inferred to be feasible based on the covariation of gene expression patterns observed in measured cells. In practice, we ignore projections *y*_*i,k*_ when the predicted cell type abundances *α*_*i,k*_ is small (e.g. <5%).

With scProjection, we achieve state-of-the art deconvolution performance in predicting cell type abundances with across multiple benchmarks^16,17^ (**Supplemental Note 1**). scProjection particularly performs well at estimating rare cell type abundances compared to other approaches. The rest of this study focuses on the projection task of inferring cell states of individual cell types.

### Projections distinguish within-cell type variation in gene expression patterns

Initially, we established scProjection’s ability to map mixed RNA samples to the correct transcriptional state for each contributing cell type. To do so, we conducted a series of simulation experiments in which a pair of cell states were selected from distinct neuron cell types, L2/3 IT and L6b, profiled in a recent human cortex cell atlas^5^. To impose a tiered difficulty, we chose these two neuronal subclasses s which are variable in their heterogeneity: L2/3 IT is highly variable with many cell states, and L6b is composed of five cell states (**Methods**). We repeatedly constructed mixed RNA samples by first selecting a random subtype, then selecting a cell state from that subtype, for each of L2/3 IT and L6b. The gene counts from this pair of randomly selected cells were added to form the final mixed RNA sample.

scProjection was then evaluated on its ability to map the mixed RNA sample back to the correct transcriptional state and subtype for each of L2/3 IT and L6b, when only provided with a cell atlas whose cells are annotated at the level of L2/3 IT and L6b (no subtype information was provided to scProjection). We found that scProjection mapped all 10,000 mixed RNA samples back to their correct subtype. Furthermore, we found that scProjection mapped the RNA samples to the correct and higher resolution cell state in 87% of the simulations, and the projected cell state was highly correlated to the original (Spearman rho=0.99, p < 2.2e-16) (**Supplementary Fig. 1**). This compares favorably to CIBERSORTx, which mapped each RNA sample back to the true subtype only 61% of the time, with an average Spearman correlation of rho = 0.68 to the original cell state. These findings are consistent with experiments performed on the CellBench gold standard benchmark data (**Supplementary Note 2**)

Having demonstrated scProjection can successfully project simulated data to the correct cell state and subtype, we designed an analogous experiment using experimentally measured RNA samples from single cell resolution assays. A recent Patch-seq study^18^ profiled 4,200 mouse visual cortical GABAergic interneurons from multiple layers of the mouse neocortex, of which the original study classified 1,818 of them as Sst inhibitory neurons, the most abundant class in the dataset. As we described above, Patch-seq RNA measurements typically contain RNA from the target neuron as well as neighboring non-neuronal cells, so the goal of our experiment was to perform projection to recover the cell state of the target neuron for each Patch-seq measurement (**Fig. 2a**). We first performed a sanity check by using scProjection to estimate the abundance of the Sst cell type within each of the 1,818 Patch-seq RNA measurements of experimentally defined Sst neurons. We found Sst was the cell type with highest abundance in 1,764 of the measurements; these results were consistent even when using two different single nucleus atlases of the brain (**Supplementary Fig. 2**). These results confirm we could accurately map Patch-seq RNA measurements to the correct cell type.

**Figure 2.**
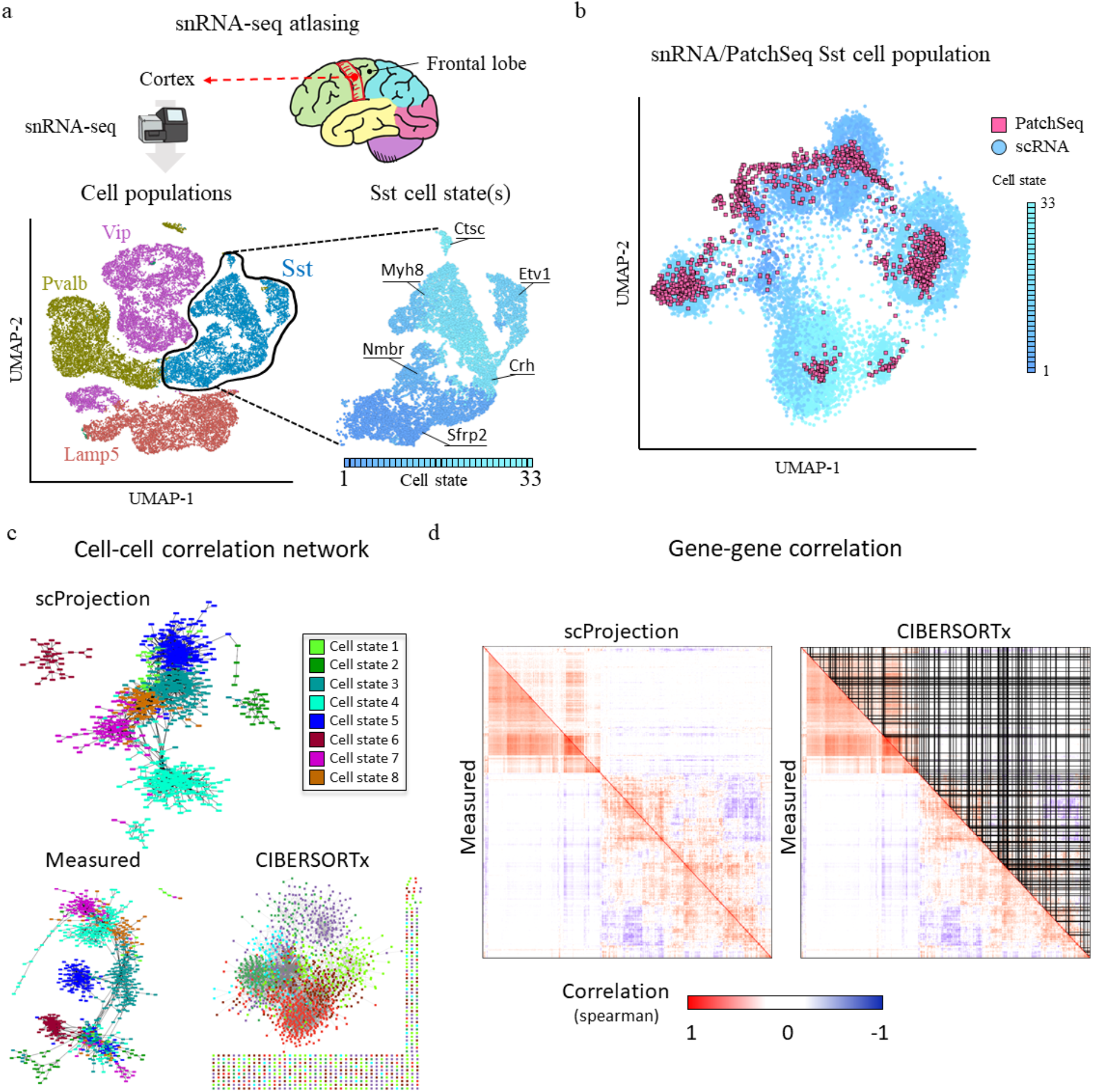
scProjection distinguishes within-cell type variation and maintains cell-cell and gene-gene network structure in Sst neurons. (**a**) Visualization of the snRNA-seq atlas of the mouse cortex used for projection of mouse Patch-seq data. We subsetted the data to four major cell types (Sst, Vip, Pvalb and Lamp5), of which Sst was further broken down into 33 distinct cell states. (**b**) tSNE plot of the measured single cell (circle) Sst neurons (from (a)) alongside the mouse PatchSeq (square) measurements projected to the Sst population. snRNA-seq cells are colored according to cell state shown in (a). (**c**) Cell-cell similarity network of the measured Patch-seq Sst cells, the scProjection-based projection of Patch-seq RNA to the Sst population, and CIBERSORTx-predicted contributions of the Sst population for comparison. (**d**) Heatmaps visualizing the gene-gene covariation patterns of the measured Patch-seq RNA (lower-triangular), versus the gene-gene covariation patterns calculated from either the projections of the Patch-seq RNA to Sst via scProjection, or the CIBERSORTx-based predictions of RNA contributions by Sst. in the upper-triangular of their respective heatmaps.

We then used scProjection to project the 1,818 Sst Patch-seq RNA measurements to an Sst single nucleus atlas^5^ (**Fig. 2a**). Because the ground-truth cell state of the Patch-seq measurements is unknown (unlike in the simulation), we instead measured the accuracy of our projections by comparing the experimentally-defined Sst subtype of the Patch-seq measurement (which is not provided to scProjection) and the known Sst subtypes of the single nucleus measurements in the atlas. In 1623 of the 1,818 neurons, the cell state of the projected Sst neurons matched the annotated cell state of neighboring neurons from the single cell atlas (Methods) (**Fig. 2b**). Similarly, we projected a separate Patch-seq dataset consisting of 45 layer 1 inhibitory neurons from two electrophysiologically-defined subclasses (SBC, eNGC) onto a broad single cell atlas of inhibitory neurons. We found the SBC and eNGC neurons were better separated after projection (Acc: 0.84) compared to the original Patch-seq RNA measurements (Acc: 0.35) (**Supplementary Fig. 3**). In total, our results on these two Patch-seq datasets suggest that scProjection distinguishes intra-cell type expression variation associated with neuronal firing patterns within the inhibitory neuron cell types

### High-fidelity maintenance of cell and gene network structure

One concern we had while designing scProjection was whether projections altered the input RNA samples as a population. That is, if two input RNA samples are similar before projection, we reasoned they should tend to be similar after projection; that is, the overall similarity structure of the input samples should remain globally consistent. On the other hand, we also would expect that the co-expression behavior of individual genes after projection would be consistent with the reference single cell atlas; genes that co-vary (and therefore are more likely to co-function) in the single cell data should also do so in the projected samples, since they represent the same cells. Therefore, to measure these population level behaviors, we constructed cell-cell and gene-gene co-expression networks before and after projection and compared them.

**Figure 2c** illustrates three inferred cell-cell co-expression networks: that of the Patch-seq measurements before and after projection to Sst, as well as from the imputed gene expression profiles of CIBERSORTx. Overall, the structure of the cell-cell network after projection more closely resembles the before-projection measured network (Jaccard: 0.72) compared to CIBERSORTx, suggesting scProjection maintains the overall structure of a set of input samples compared to CIBERSORTx (Jaccard: 0.21). Similarly, **Figure 2d** qualitatively compares the inferred gene co-expression network of the measured Sst scRNA-seq data, to both the projected samples from scProjection, as well as the imputed samples from CIBERSORTx. scProjection’s network more closely resembles the measuredSst co-expression networks, in comparison to CIBERSORT which fails to impute many genes as visualized by the black lines.

### Detection of novel spatial expression patterns of enterocytes in the intestinal epithelium

We envisioned that one primary application of scProjection is to infer single cell transcriptomes from RNA measurements produced by spatial transcriptome technologies, in order to detect spatial gene expression patterns in tissues. Technologies such as Slide-seq^8^, LCM-seq^9^ and Visium by 10x Genomics capture RNA from different spots of a tissue slice. Each spot potentially contains RNA contributions from more than one cell in close proximity (**Fig. 1a**). Therefore, the RNA from each spot can be viewed as a miniature bulk RNA sample composed of a small number of cells, from which we want to extract single cell transcriptomes for each contributing cell type through projection.

We initially analyzed a dataset collected by Moor et al.^19^ in which they performed LCM-seq on five distinct regions, or zones, of the intestinal villus, as well as separately collected a scRNA-seq cell atlas from replicate intestinal villi. They identified spatial expression patterns in the dominant cell type, enterocytes, by (1) identifying marker (landmark) genes for each zone using the LCM-seq data, (2) assigning zone labels to the scRNA-seq cells using landmark genes, and (3) predicting zone-specific expression through zone-specific averaging of the labeled scRNA-seq data. We reasoned that identification of landmark genes from LCM-seq data could be difficult since LCM-seq captures contributions from multiple cell types, thus yielding poor labeling of the single cell atlas cells. We therefore avoided this critical landmark gene selection by taking the opposite approach: we use scProjection to project the zone-specific LCM-seq samples to the enterocyte single cell atlas, in order to extract the enterocyte expression patterns within each zone. This approach would explicitly disregard contributions of non-enterocytes to each LCM-seq sample.

**Figure 3a** illustrates the projections of the LCM-seq data to the enterocyte single cell atlas, where the single cells are labeled according to Moor et al^19^. The LCM-enterocyte projections are generally proximal to the single cells assigned to the same zone by Moor et al., suggesting our approach is overall consistent with that of Moor et al. However, our approach identifies 3-fold more zone-specific spatial expression patterns compared to the genes identified by the Moor et al (**Fig. 3b**). To validate the predicted enterocyte zone-specific expression patterns, we compared our predicted zone expression patterns to the smFISH expression quantifications and the original LCM-seq measurements provided in the original study. We found that across a small set of validated landmark genes (Ada, Slc2a2, Reg1), our spatial expression predictions followed with the smFISH quantifications performed in Moor et al. (**Fig. 3c**). Furthermore, our approach identified zonation patterns in genes such as Pkib, Slc2a13 and Fam120c which were not identified by the Moor et al. spatial reconstruction approach yet are clearly zone-specific according to the original LCM-seq experiments (**Fig. 3c**). These results in total suggest RNA projections improve our ability to identify zone-specific expression patterns in dominant cell types such as the enterocytes.

**Figure 3.**
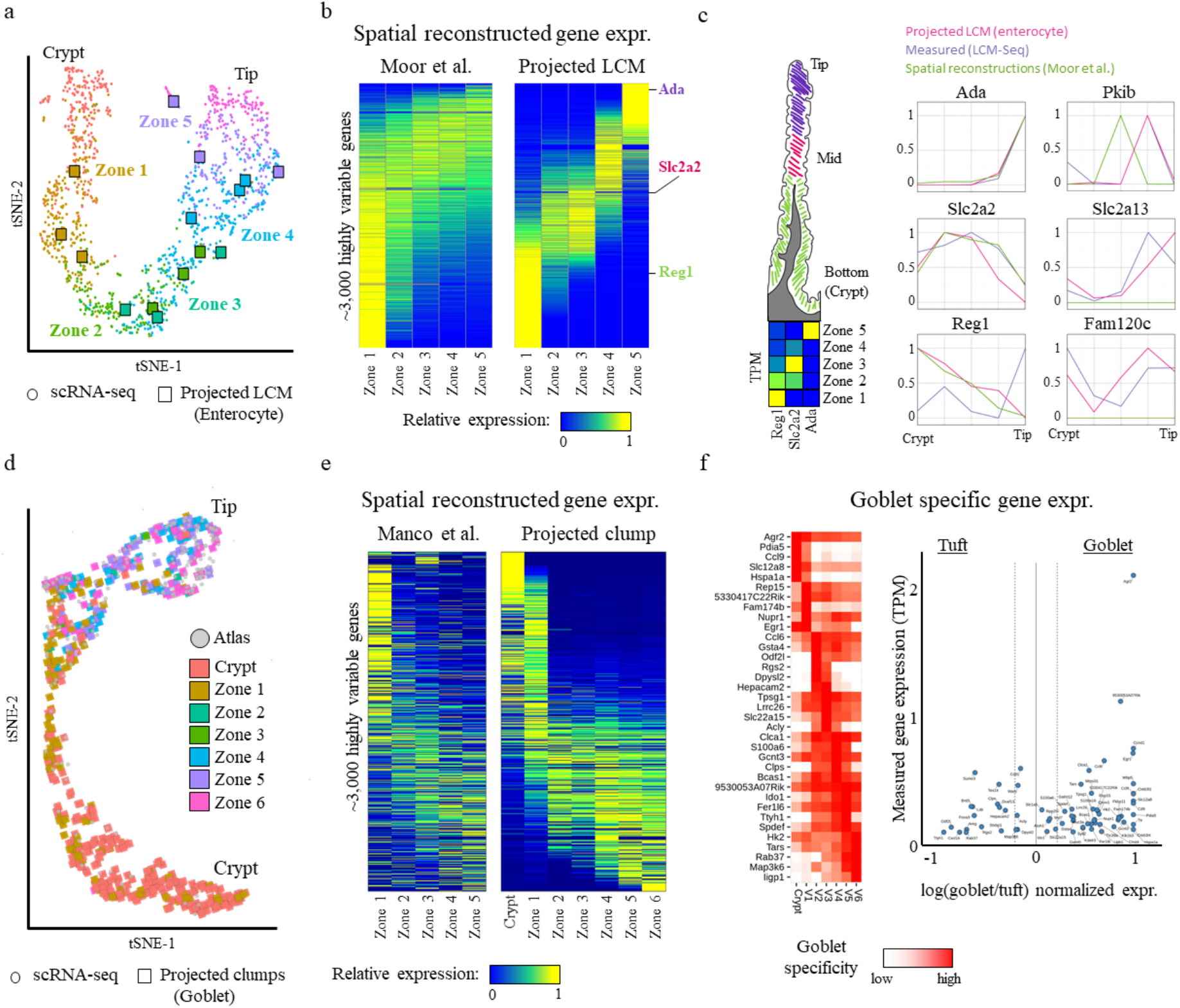
Projection refines spatial expression patterns in common and rare cell types of the intestinal villus. (**a**) tSNE plot of the single cell atlas (circles) and projected LCM samples (squares) across the zones of the intestinal villus. Single cells are colored based on their zone assignment by Moor et al. (**b**) Heatmap visualizing the spatial expression patterns of the top 3,000 highly variable genes using the spatial inference approach of Moor et al. on the left and after projecting the LCM samples with scProjection on the right. Three marker genes (rows) are labeled: Ada, Slc2a2 and Reg1. On the right is a schematic of a single intestinal villus, along with the expected dominant zone of expression for Ada, Slc2a2 and Reg1. Shown below the villus is the actual measured expression pattern of Ada, Slc2a2 and Reg1 in the LCM data of the five zones. (**c**) Line plots comparing the measured and projected expression of top zonated genes across the intestinal villus. (**d**) tSNE plot of the single cell atlas (circles) and projected clump-seq (squares) as annotated by the enterocyte component within each clump. (**e**) Heatmap visualizing the spatial expression patterns of the top 3,000 highly variable genes in the goblet containing clump-seq samples using the approach of Manco et al. on the left and after projecting with scProjection on the right. (**f**) Heatmap visualizing the expression of the union of the top 5 zonated genes per zone in the goblet containing clumps. The scatter plot on the right visualizes the divergence in expression of zonated genes between goblet and tuft containing clumps

### Rare cell types of the intestinal villus can be spatially resolved

Projection of an RNA sample onto the single cell atlas of a target cell type intuitively requires sufficient abundance of the target cell type within the RNA sample in order to be successful. scProjection predicted enterocytes to contribute 90% of the LCM-seq RNA on average. In contrast, populations such as the secretory (goblets, tuft) cells are rare: for example, goblets only contribute 8% of the LCM-seq RNA on average^20^, while tuft cells are only contribute 1% of the LCM-seq RNA on average (**Supplementary Fig. 4**). The mucus-producing goblet cells^21,22^ and chemosensory tuft cells^23^ play an important role in the protection of cells in the intestinal villus as well as communication with other stroma cell types^24^. Manco et al.^20^ captured these rare cell types in the intestinal villi by performing RNA-seq on clumps of physically-adjacent cells in the intestinal villus, through incomplete dissociation of the tissue. Because of the high abundance of enterocytes and rare occurrence of goblet and tuft cells, most clumps will contain primarily enterocytes, and only occasionally contain goblet or tuft cells. To derive spatial expression patterns of the rare cell types, Manco et al. predicted the zone of the entire clump by comparing clump expression against a spatial reference from the Moor et al.^19^ work described above, then assigned that zone label of the entire clump to the secretory cells within the same clump. We hypothesized that by replacing the zone-prediction step in Manco et al. with our projection approach used above for the enterocytes, we can further identify goblet and tuft specific spatial patterns of expression across the intestinal villus.

Our general strategy was to first train the scProjection VAE components on individual cell types within a single cell atlas of the intestinal epithelium, which captured enterocytes and rare secretory types including goblet and tuft cells^20,25^. We then simultaneously project each clump to the enterocyte cell type and the secretory cell types (goblet or tuft) separately. We predicted the zone of the entire clump based on the zone-specific LCM-enterocyte projections similar to above (see Methods). We computed zone-specific expression patterns of goblet (or tuft) cells by averaging clump-goblet (or clump-tuft) projections that were predicted to land in the same zone.

We focused first on the mucus-producing goblet cells, because while rare, there were more goblet-containing clumps available to robustly estimate zone-specific expression compared to tuft cells. From an initial set of 6,824 clumps, we identified 1,084 clumps that contained at least 40% cell type abundance from goblet cells. From the 1,084 goblet-containing clumps, we projected these clumps to the goblet single cell population (n=314) to identify spatial gene expression patterns. **Figure 3d** illustrates the 1,084 clumps projected onto the goblet single cell atlas, where the clumps and single cells are labeled according to Manco et al^20^ (**Supplementary Fig. 4**, Methods). The projected clumps were generally proximal to the single cells assigned to the same zone by Manco et al., suggesting our approach generally consistently captures zone-specific gene expression. Using our projections of the clumps, we predicted 2480 genes that exhibit zone-specific goblet expression patterns, compared to 972 zone-specific genes identified by Manco et al.’s approach (**Fig. 3e**). To validate the predicted goblet zone-specific expression patterns, we compared our 2480 zone-specific genes with goblet specific landmark and mucus associated genes (Methods) whose tendency for villus-tip expression was identified in Manco et al. We found that our spatial expression predictions from the clumps were followed reported zonated expression and smFISH quantifications (**Supplementary Fig. 5**) suggesting our projections can accurately capture zone-specific expression of goblet cells.

As members of the secretory cell class, the goblet and tuft cells derive from a common progenitor^23^, and have previously been noted to both express common immune modulatory pathways^23^. We therefore wondered whether we could identify genes that are both zone-specific, and specific to a single lineage (goblet or tuft). We therefore identified clumps that contained at least 40% cell type abundance from the tuft cells, then projected those clumps to the tuft cell population to identify tuft zone-specific expression patterns similarly to the goblet analysis above (**Fig 3e, Supplementary Fig. 4**). To identify goblet (or tuft)-specific, zone-specific expression patterns, we computed the ratio of goblet and tuft specific expression for each gene per zone, and identified the top five genes per zone exhibiting goblet specific expression (log(goblet/tuft) > 0.9) (**Fig. 3f**). The goblet-specific gene, Agr2, in the crypt zone stands out as highly expressed and goblet specific (**Fig. 3f**), and is a known landmark^26^. However, most genes that were specific to goblet or tuft were expressed at relatively low levels (TPM<1), suggesting the differences in expression between goblet and tuft may be driven by noise.

### Transcriptome imputation helps infer global spatial expression patterns in the brain

Imaging-based spatial transcriptome technologies such as MERFISH and seqFISH enable imaging of individual transcripts in 2D tissue slices and therefore provide insight into spatial expression patterns at sub-cellular resolution. However, these technologies have two drawbacks: (1) it may not be practical to spatially profile all genes in the genome; for example, MERFISH experiments have profiled only hundreds of transcripts^27^ to date, and (2) imaging pipelines^28^ are required to segment the images into cells in order to compute single cell expression patterns, which can be an error-prone process and lead to transcripts from adjacent cells being grouped into one ‘cell’^28^.

To address the limitation of the smaller number of genes that can be measured by imaging-based technologies such as MERFISH and seqFISH, we modified scProjection so that even with a small, refined set of measured genes for the input RNA samples, scProjection would project those RNA samples to genome-wide expression profiles of individual cell types. Intuitively, scProjection uses direct and indirect correlation between the measured genes and missing genes (assessed from the single cell atlas) to perform non-linear imputation of gene expression measurements. In that way, scProjection could be used to simultaneously attain single cell expression measurements and impute the rest of the genome’s expression signal.

In a study of neurons from the hypothalamic preoptic region of the mouse brain, Moffit et al. assayed 155 marker genes across millions of neurons using MERFISH and generated a matched scRNA-seq cell atlas. Using scProjection, we imputed genome-wide expression patterns for the entire MERFISH dataset spatially profiling millions of neurons. Labeling each MERFISH sample by the cell type that contributes that most RNA, we found scProjection recovered the spatial organization of Oligodendrocytes across slices from the mouse brain defined by Bregma indices (**Fig. 4a**). More specifically, the oligodendrocytes spatially organize into one cluster at Brega 0.26, then eventually diverge into two populations by Bregma -0.29. To explore potential functional implications of the segmentation of oligodendrocytes from one into two spatial regions, we computed Bregma index-specific expression patterns of Oligodendrocytes between Bregma 0.26 and -0.29 and identified many genes with clear differential expression patterns across the two distal Bregma indices (**Fig. 4b**). Of particular note are Calca and Dpp10, both of whom are associated with oligodendrocyte differentiation that occurs along the bregma axis with immature and mature oligodendrocytes occupying separate compartments of the hypothalamus^27^. Neither of these markers belonged to the 155 marker gene set measured by MERFISH in the original study. scProjection therefore helps identify genes with spatially distinct expression patterns, even if they were not measured in the original spatial transcriptome assay.

**Figure 4.**
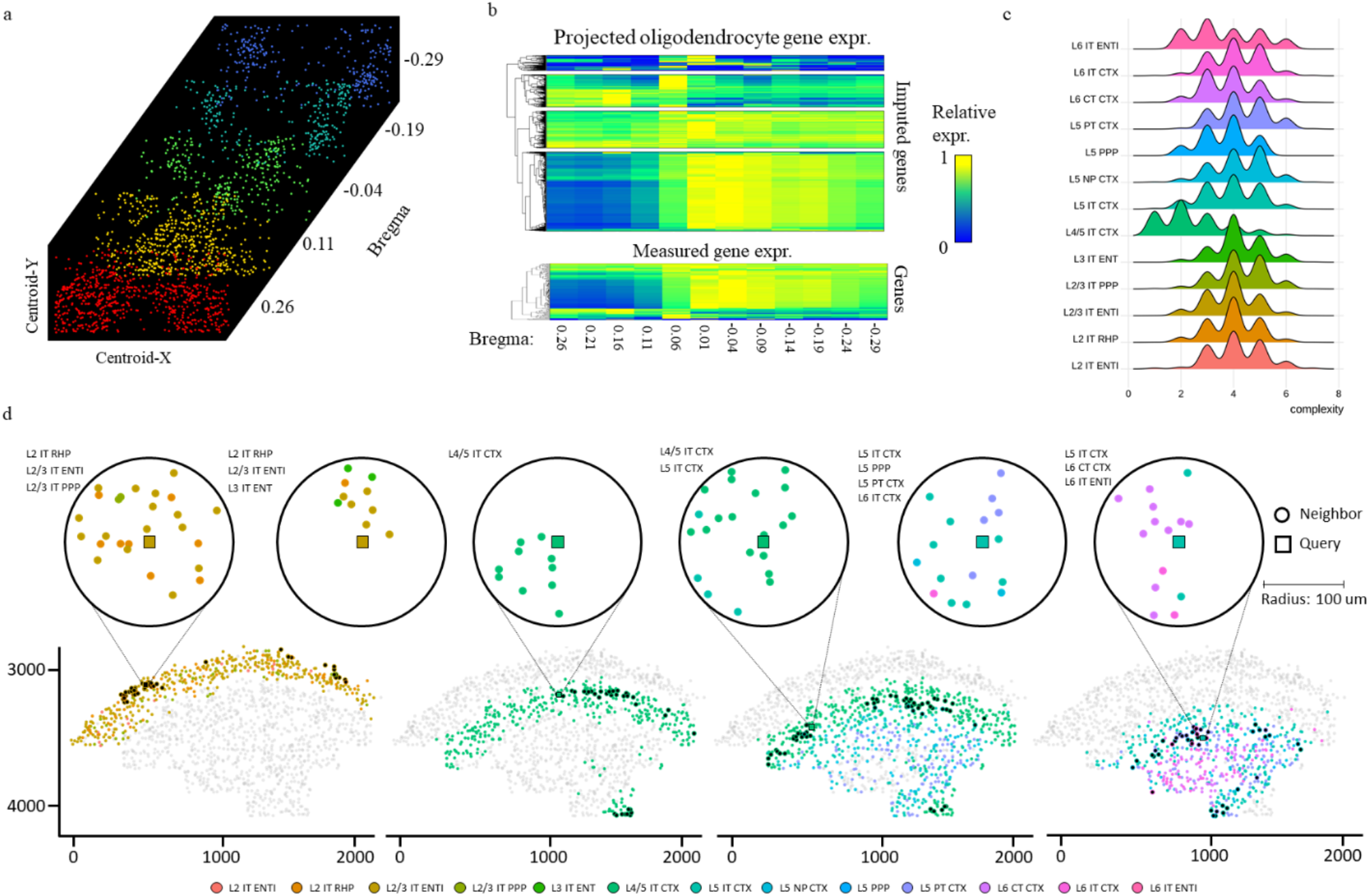
Imputation and high-resolution label transfer identifies spatial expression patterns in the brain. (**a**) Stacked tSNE plots of oligodendrocyte populations identified according to dominant cell type with scProjection across Bregma indices from Moffit et al. (**b**) Heatmap visualizing the spatial expression patterns within the oligodendrocytes of imputed (top) and measured (bottom) genes from the original study. (**c**) Neighborhood density plots for each cell type annotated by scProjection, where the x-axis indicates the neighborhood complexity for each cell. (**d**) tSNE plot of a single slice separated by layer type of the neurons according to the post significant neighborhoods highlighted in the circle plots for a few neurons from the mouse cortex mapped by Zhang et al. and as annotated by scProjection.

### Identification of spatial motifs in the primary motor cortex

The identification of spatial gene expression patterns is a task often performed at the individual gene level; many approaches have been developed to identify non-random spatial single gene expression patterns ^29,30^. Spatial patterning within tissues extends well beyond the level of individual gene expression patterns, however. At a coarse level, the mammalian brain organizes neurons into functional neighborhoods that vary with cortical depth^12,31^. Interneurons from different layers of the cortex are widely recognized as distinct in their transcriptome and function ^1,5,18^. We hypothesized that there might be more localized structure to cell type organization in the brain, involving potentially small groups or types of cells that frequently spatially co-occur together. We term these larger groups of co-occurring cells “spatial motifs”.

To identify spatial motifs as a function of cortical depth, we analyzed data from a recent MERFISH study by Zhang et al.^31^ in conjunction with a million-neuron atlas from Yao et al.^5^ of the mouse primary motor cortex (MOp). We used scProjection to infer a revised high resolution cell type label for each MERFISH measurement by projecting MERFISH measurements to the snRNA-seq atlas and assigning discrete labels based on the taxonomy of Yao et al., which defines 129 cell types what broadly fall under the category of glutamatergic, GABAergic, and non-neuronal subtypes.

We first performed neighborhood analysis by quantifying, for each high-resolution label, the complexity of its physical neighborhood within a 100um radius. More specifically, we define the complexity of a cell’s neighborhood as the number of distinct cell types present in a 100um radius of the cell (Methods). For each brain slice, we computed the distribution of neighborhood complexities of glutamatergic (excitatory) neurons as a function of cortical depth and high-resolution cell type annotated by scProjection. Comparing the neighborhood complexity of excitatory neurons across cortical depth revealed that most cortical depths were comparably complex (mean complexity: 4 cell types), with the notable exception of L4/5 IT CTX neurons which were overall less complex (mean complexity: 1.5 cell types) (**Fig. 4c**). 24% of the L4/5 neuron cells had homogenous neighborhoods that contained no neurons from any other layer, an observation unique to the L4/5 neuron cells.

Having shown that mapping MERFISH samples to high resolution cell types can enable the identification of diverse neighborhood types, we next looked for the existence of spatial motifs, defined as spatial neighborhoods consisting of a specific set of cell types that are unlikely to occur by chance. We assigned each cell into a spatial neighborhood type based on the number of cell types within a 100um distance. We then counted the number of cells assigned to each spatial neighborhood type, and permuted the cell type labels 1,000,000 times, making sure cell labels only permute within cells of the same layer. Through comparison with simulated neighborhood occurrences, we identified a diverse set of 19 significant (permuted p < 0.05/1000000, n>50) neighborhood types ranging from homogenous L4/5 populations to neighborhoods which exists on the L2/3 and L6 boundaries (**Figure 4d**). Many of these neighborhoods involved cell types from multiple layers, even though our permutations kept cell labels of the same layer together. This suggests non-random placement of cell types near layer boundaries. These spatial motifs occurred frequently; on average, 231 cells were assigned to each of the 19 significant spatial motifs. Of note, the L4/5 IT CTX neurons were the only high-resolution cell type to form islands of neurons containing only the same type (Complexity: 1) within 100um. By annotating higher resolution high-resolution cell type annotations onto the MERFISH data with scProjection we can uncover neighborhood structure underlying coarser cell type spatial variation.

### Projection of Patch-seq RNA improves identification of connections between gene expression and neuron electrophysiology

Besides spatial transcriptome technologies, there are several other single cell resolution assays that could benefit from scProjection. For example, Patch-seq^10^ is a protocol for jointly measuring the RNA, electrophysiological (ephys) and morphological properties of individual neurons, and is critical for linking the molecular and cellular properties of neurons. Patch-seq uses a micropipette to puncture a neuron in order to simultaneously measure its RNA and electrophysiological properties. When applied to *in vivo* or *ex vivo* slices of brain tissue, the micropipette passes through other surrounding cells in order to reach the neuron of interest, leading to the RNA measurements containing contributions from both the target neuron as well as surrounding glial cells^11^. scProjection analysis of several Patch-seq studies indicates cell type abundances from non-neuronal cells are predicted to be as high as 30%, suggesting significant contamination of RNA (**Fig. 5a**). We therefore hypothesized that projecting Patch-seq RNA measurements to a single cell atlas of neurons would reduce the effect of contaminating RNA and improve downstream analyses such as correlating gene expression measurements to electrophysiological measurements of neurons.

**Figure 5.**
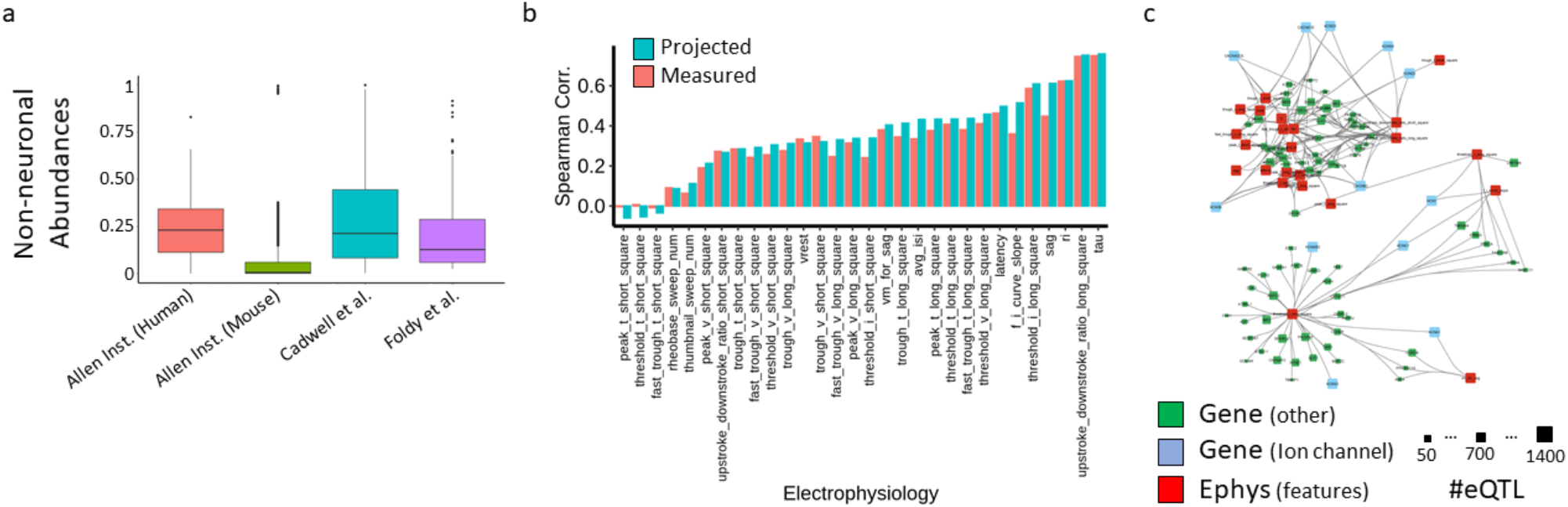
Projection of Patch-seq RNA links molecular measurements to electrophysiology of neurons. (**a**) Box and whisker plots visualizing the abundances of non-neuronal RNA estimated by scProjection across all samples of multiple PatchSeq studies. (**b**) Bar plot of the accuracy (based on Spearman correlation) of gene expression-based prediction of electrophysiology measurements, when predictions are made using either the original measured RNA, or the Sst projected PatchSeq samples. (**c**) Gene – electrophysiology correlation network, where edges are between significantly correlated genes and electrophysiology features. Node size is proportional to the number of eQTLs identified in the xQTL study of the ROSMAP cohort.

We applied scProjection to a set of 4,200 Patch-seq measurements targeting mouse GABAergic neurons^18^, together with a reference atlas of the mouse brain^5^. Of the 4,200 measurements, scProjection predicted that 1,912 of them were primarily targeting Sst inhibitory neurons (**Supplementary Fig. 6**), consistent with the fact that these 1,912 assayed neurons were experimentally identified as Sst before Patch-seq. We focused our experiments on the 1,912 predicted Sst inhibitory neurons because they were the best represented type of neuron, and therefore projected the 1,912 Patch-seq measurements to the Sst single cells sequenced in the reference atlas.

Here we assumed that more accurate Patch-seq RNA measurements should enable better prediction of ephys properties of neurons from gene expression levels. To this end, our RNA projection enabled a mean increase of 27% prediction accuracy of two ephys features, sag and latency, from genome-wide expression profiles (spearman correlation of 0.62 compared to 0.43, p = 5e-18, rank sum test) (**Fig. 5b**), while other features were comparable before and after projection. Additionally, we found that our RNA projections identify significant (q<0.05) cell type-specific correlations in Sst projected ion channel gene expression and ephys properties (**Supplementary Fig. 7**). These results together suggest that RNA projections remove noise driven by the presence of non-neuronal abundances, which leads to better identification of connections between gene expression and neuron electrophysiology.

Having used scProjection to establish more gene-ephys connections than could be previously appreciated from the original Patch-seq data, we further hypothesized that genetic variation may drive systematic changes in some ephys features, through changes in gene expression patterns. We extracted cis-eQTLs detected in the human dorsolateral prefrontal cortex from the ROSMAP consortia^4^, and found that 91 genes’ expression levels were both associated with genetic variation, and also correlated with ephys features of neurons. Although gene-ephys connections were identified via correlative analysis and so we cannot directly infer that these eQTLs will causally influence ephys properties in general, we looked specifically at ion channel genes because they play critical roles in establishing ephys responses to neuron stimuli. We found 12 ion channels associated with neuronal firing and under genetic control, of which 58% of them were only identified after projection (but not with the original Patch-seq measurement). We also identified 79 genes not annotated as ion channels that are also associated with electrophysiology and eQTLs (**Fig. 5c**). In fact, 83% of all genes associated with the 31 ephys features are not ion channel genes. While much of the focus of interactions between genes and electrophysiology is on ion channels, our results suggest there may be many more genes that either directly influence ephys in novel ways, or indirectly interact with ion channels for example.

## DISCUSSION

In our experiments, we have demonstrated the utility of projections for the analysis of diverse single cell resolution assays such as spatial transcriptomes and Patch-seq. At its heart, projection maps RNA samples into the cell state space defined by a single cell atlas. Therefore, RNA projections can also potentially play a role in up-sampling the per-cell sequencing depth of spatial and multi-modal sequencing assays, by projecting lower depth samples into a high depth cell atlas. For example, because RNA capture is not per-cell but per-spot for technologies such as Slide-seq, the number of effective transcripts sequenced can vary spot to spot^8^. Furthermore, mRNA capture efficiencies can vary between protocols^32^, and technologies such as SMART-seqv2 yield significantly high read depth per cell compared to 3’ tagging technologies such as the 10x Chromium ^33^. scProjection can be used to project RNA samples sequenced from specialized spatial and multi-modal sequencing assays into a deeply sequenced scRNA-seq atlas for example, in order to increase the resolution of the resulting gene expression profiles. This is conceptually similar to the process of imputation that we demonstrated in our MERFISH results, though imputation is typically cast as a problem of filling in zero transcript counts rather than up sampling both non-zero and zero counts.

RNA projections are complementary to deconvolution methods. The goal of deconvolution methods^34–37^ is primarily to estimate the cell type abundances of a set of reference cell populations within a single RNA sample, and is a very well-studied problem dating back several decades^38^. While scProjection also computes such cell type abundance to a set of populations, its primary goal is to distinguish intra-cell type variation by also mapping the RNA sample onto the precise cell state within each of the cell type populations that best represents the expression profile of those cell types within the RNA sample. scProjection therefore distinguishes intra-cell type variation, whereas deconvolution methods principally focus on differences in cell type abundances in a sample.

A major feature of scProjection is that it implicitly fits a probability density function (PDF) over the cell state space for each cell type. This is advantageous for several reasons. First, this enables scProjection to reason about the relative frequency of a cell state observed in the training data, where more frequently observed states have higher probability of being projected to. Second, it enables scProjection to interpolate between observed cell states when the training data is small, which can be important for training on rare cell types or on data from smaller studies. Third, scProjection can also naturally ignore outlier sequenced cells in the training data because they will not appear often in the cell atlas. In contrast, a number of other methods either average the expression profiles all cells of the same type such as CIBERSORTx that we tested here^34^, or only map RNA samples to measured single cells in the atlas^39^. Methods that average cells of the same type together will be sensitive to outliers, and more importantly will be unable to account for variation within a given cell type.

One of the caveats of scProjection and related methods, is that by projecting RNA measurements to a reference single cell atlas, scProjection assumes that the single cell atlas contains accurate representations of the cell state of cell populations within the RNA sample. There could be scenarios where this is false; for example, projecting RNA from a spatial transcriptome assay of (liver) hepatocellular carcinoma samples to a normal liver atlas would miss expression variation in hepatocytes that is driven by carcinomas. Therefore, if no suitable single cell atlases are publicly available, it would make sense to collect scRNA-seq data on some biological replicate samples in addition to the spatial transcriptome datasets. This experimental design of collecting both scRNA-seq as well as spatial transcriptome data is common^8,19,40,41^ so we expect this caveat to not limit the widespread applicability of scProjection.

Finally, we envision applications of RNA projections beyond what we have illustrated here. For example, databases such as the Gene Expression Omnibus (GEO) catalog gene expression data from bulk RNA samples collected since RNA sequencing was first deployed. Using the increasing number of single cell atlases derived for different tissues and cell types across organisms, scProjection can be used to re-analyze historic bulk RNA samples to extract average cell states for individual cell populations that contribute to the bulk RNA sample. Cell type-specific changes in case-control studies could then be inferred, as could cell type-specific eQTLs from genetic studies of disease, for example.

## ACKNOWLEDGEMENTS

GQ was supported by NSF CAREER award 1846559. This project has been made possible in part by grant number 2019-002429 from the Chan Zuckerberg Foundation. This work was supported by the National Institute of Child Health and Human Development P50 HD103526.

## METHODS

### scProjection overview

Our framework, scProjection, projects *N* gene expression profiles ***b***_*n*_ ∈ *B* generated from RNA samples into each of *K* different cell populations represented in a reference single cell atlas, yielding a new set of gene expression profiles ***x***_*n,k*_, for *k* = 1, …, *K*. scProjection also estimates *α*_*n,k*_, the proportion of RNA contributed by each cell population *k* to sample *n* (**Fig. 1**). scProjection assumes that each ***b***_*n*_ is a weighted linear combination of the cell population-specific projections ***x***_*n,k*_:

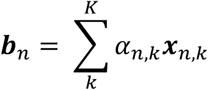

Only ***b***_*n*_ is formally observed, and the goal is to estimate *α*_*n,k*_ and ***x***_*n,k*_.

To perform estimation, scProjection leverages a separate reference single cell atlas in which single cells ***s***_*j,k*_ (representing the *j*^th^ cell sequenced for cell population *k* in the atlas *S*) have been sequenced. In the first step, scProjection trains a deep variational autoencoder (VAE) separately for each cell population *k* using all single cells sequenced for cell population *k* (*s*_*,*k*_), yielding a parameter set {*ϕ*_*k*_, *θ*_*k*_} (representing the encoder and decoder parameters, respectively) for each cell population *k*. After training, each VAE implicitly defines the set of cell states that projections into cell population *k* (*x*_*n,k*_) can occupy. In the second step, the VAEs with trained parameters 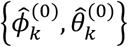 are used to get initial projections 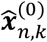 by inputting each ***b***_*n*_ into the *k*^th^ VAE and sampling from the output to estimate 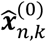. In the second step, we estimate the RNA proportions 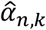 by solving the above equation by using linear regression by setting 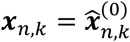. Finally in the third step, we fix the mixing proportions 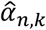, and re-update all VAE parameters {*ϕ*_*k*_, *θ*_*k*_} simultaneously to improve estimates of ***x***_*n,k*_ by maximizing the reconstruction of each ***b***_*n*_.

### scProjection training of cell population-specific VAEs (Step 1)

scProjection uses VAEs to perform the projection of RNA samples ***b***_*n*_ into the gene expression space of each cell population *k* to yield the projection ***x***_*n,k*_. The set of cell population-specific VAEs are identical in network structure and are comprised of a deep encoder network parameterized by weights *ϕ*_*k*_, and decoder network parameterized by weights *θ*_*k*_. To train the VAEs, we optimize the following objection function with respect to the VAE parameters {*ϕ*_*k*_, *θ*_*k*_}:

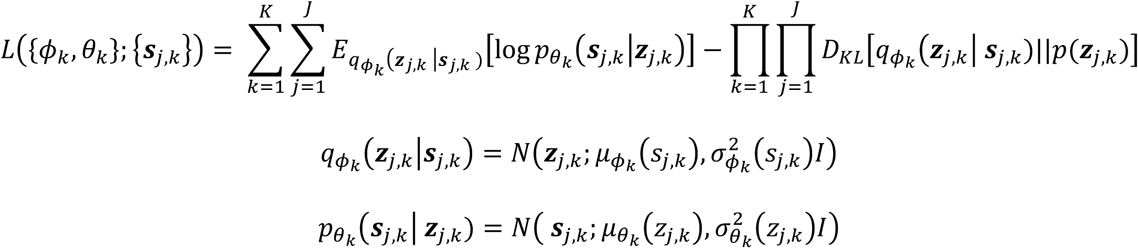

The functions 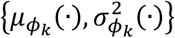 and 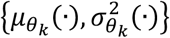 represent the mean and variance of the normal distribution predicted by the encoder and decoder, respectively. The parameters of the VAEs {*ϕ*_*k*_, *θ*_*k*_} are regularized through 30% dropout [13], batch normalization [14] and L2 weight regularization to ensure robust training. ADAM [15] is used for optimization with a decaying learning rate starting at 1e-3 and a smooth warmup of the KL term in the ELBO, which has been shown to produce more accurate reconstructions ^42^. We denote the trained VAE parameters By 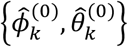.

For the experiments in which we impute genome-wide expression measurements from limited sets of marker genes such as those measured by MERFISH, the structure of the VAE becomes asymmetric with the input measurements to the encoder defined by a subset of gene expression measurements *G*_*e*_ ⊆ *G* (corresponding to marker genes). The decoder output is still defined by the full set of gene expression measurements *G* made in the single cell atlas. Only estimates of those genes *G*_*e*_ directly measured in mixture samples *b*_*n*_ are used in subsequent steps of scProjection.

### scProjection estimation of cell type abudnacne of each cell population (Step 2)

Here, scProjection projects each RNA sample ***b***_*n*_ to each cell population *k* via the VAE parameterized by 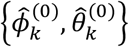 to estimate 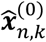:

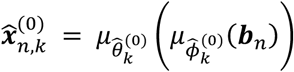

Then, we estimate the mixture proportions *α*_*n,k*_ and nuisance parameters of a multi-layer perceptron 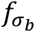 (and hold all other variables fixed) by optimizing the following objective function:

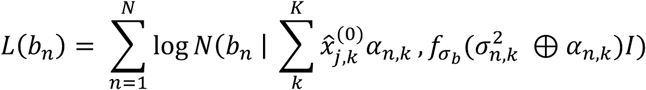

Optimization is performed with ADAM [15] and a learning rate of 1e-3 until convergence. The estimated mixing proportions 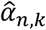 are kept fixed for the remainder of the training procedure.

### scProjection final estimates of RNA projections (Step 3)

In this step, scProjection re-optimizes the encoder and decoders of the individual VAEs {*ϕ*_*k*_, *θ*_*k*_} by minimizing the following composite objective function, which includes the likelihood of both the single cell atlas data *s*_*j,k*_ and the RNA samples *b*_*n*_:

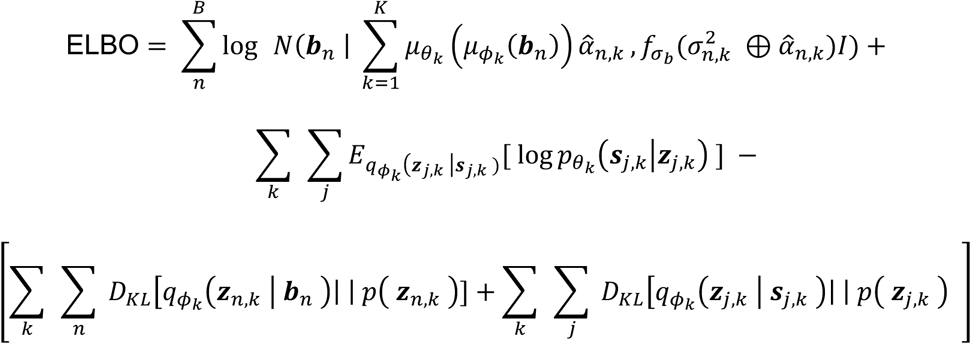

Note in this case, the VAE parameters are initially set to 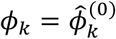 and 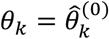 before optimization, and the parameters of 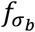 are fixed at their values estimated at Step 2. Intuitively, we are adjusting the RNA projections 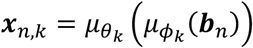 to better predict the RNA sample ***b***_*n*_, because the single cell reference data may be collected in a different experiment from the RNA samples. The single cell data are included in the objective function and serve as a regularization term to ensure identifiability of each VAE as specific to one cell population *k*. After training to obtain final VAE parameter estimates 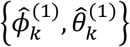, we estimate our final RNA projections 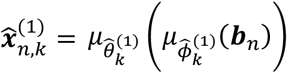.

### Acquisition and preprocessing of the intestinal villus dataset

We obtained the gene expression matrices for the LCM-seq, scRNA-seq and spatial reconstructions experiments described in Moor et al.^19^ from GSE109413 and https://doi.org/10.5281/zenodo.1320734. We independently normalized the count matrices to TP10K, then scaled and centered using Seurat’s NormalizeData (without log transform) and ScaleData functions. We retained the union of the marker genes of each cell type identified from the original study, together with the top 2,000 variable genes of both LCM-seq and scRNA-seq.

### Acquisition and preprocessing of the brain MERFISH dataset

We obtained the processed MERFISH gene luminescence matrix described in Moffitt et al.^27^ from dryad.8t8s248 and the scRNA-seq count matrix from GSE113576. We independently preprocessed each data modality by normalizing to TP10K, then scaled and centered using Seurat’s NormalizeData and ScaleData functions. We removed entire cell types from the scRNA-seq data that had no analog in the MERFISH experiments and are defined in Table S9. We retained the union of the marker genes of each cell type identified in the original study, together with the top 2,000 variable genes across the entire scRNA-seq atlas.

### Acquisition and preprocessing of the mouse Patch-seq dataset

We obtained the gene count matrix for the mouse Patch-seq experiments described in Berg et al.^43^ from portal.brain-map.org/explore/classes/multimodal-characterization on January 2019. We discarded samples that did not pass QC as defined in the original paper in both the RNA and electrophysiology modalities. We normalized the count matrix to TP10K (without log transform), then scaled and centered using Seurat’s NormalizeData and ScaleData functions. We retained the union of the marker genes of each cell type identified from the original study, together with the top 2,000 variable genes across each of the cell types defined in the snRNA-seq.

### Acquisition and preprocessing of the mouse brain atlas

We obtained the gene count matrix for the human brain atlas described in Yoa et al.^5^ from the Allen Institute Cell Types database: RNA-Seq data page on the Allen Institute’s webpage. We normalized the count matrix to TP10K, then scaled and centered using Seurat’s NormalizeData and ScaleData functions. We retained the union of the marker genes of each cell type reported in the original study, together with the top 2,000 variable genes across each of the cell types defined in the snRNA-seq. Sublcass annotations were provided for each cell and were used to filter out L2/3 and L6 for validation experiments

### Acquisition and preprocessing of the Tasic et al. mouse brain atlas

We obtained the gene count matrix for the mouse brain atlas described in Tasic et al. from the Allen Institute Cell Types database: RNA-Seq data page on the Allen Institute’s webpage. We normalized the count matrix to TP10K, then scaled and centered using Seurat’s NormalizeData and ScaleData functions. We retained the union of the marker genes of each cell type reported in the original study, together with the top 2,000 variable genes across each of the cell types defined in the snRNA-seq.

### Acquisition and preprocessing of the CellBench benchmark

We obtained the gene count matrix for the RNA mixture experiments in CellBench described in Tian et al.^44^ from the R data file mRNAmix_qc.RData available on GitHub (https://github.com/Shians/CellBench). We normalized the count matrix to TP10K (without log transform), then scaled and centered using Seurat’s NormalizeData and ScaleData functions. We retained the union of the marker genes of each cell type identified by CIBERSORTx, together with the top 3,000 variable genes computed separately on the RNA mixtures profiled on CEL-Seq2 and SORT-Seq.

### Acquisition and preprocessing of the ROSMAP-IHC benchmark

We obtained the gene count matrix for the bulk-RNA experiments and IHC measurements described in Patrick et al. from the R data files available on Github (https://github.com/ellispatrick/CortexCellDeconv). We normalized the count matrix to TP10K (without log transform), then scaled and centered using Seurat’s NormalizeData and ScaleData functions. We retained the union of the marker genes of each cell type reported in Darmanis et al.^45^, together with the top 2,000 variable genes.

### Execution of deconvolution methods

In the two sections below on benchmarking cell proportion estimations in different datasets, we compared scProjection against CIBERSORTx^34^, MuSiC^35^, NNLS, dtangle^36^, DSA^37^, and single gene deconvolution. Each method was run based on method-specific guidelines provided by the original authors and following the workflows defined by in tutorials for each approach. Prior to running each method, the FindVariableGenes function implemented in Seurat was used to identify the most variable genes for a consistent subsetting of the data matrices. CIBERSORTx was provided counts for all highly variable genes in the scRNA-seq data along with cell type annotations to create a signature matrix. Then counts for all highly variable genes in the mixture data were provided to CIBERSORTx which then estimates RNA proportions. MuSiC was provided counts for all highly variable genes in the scRNA-seq and mixture data along with cell type annotations. NNLS (as implemented by us in R) was provided the TPM values for all highly variable genes in the scRNA-seq and mixture data. Proportions from NNLS for cell type *k* were computed by summing the learned weights across all cells annotated as cell type *k*; this was repeated for each cell type and each mixture sample. dtangle was provided with a mean count vector per cell type in the scRNA-seq data and the original counts from the mixture data along with cell type markers and annotations. DSA was provided with the original counts for the mixture data and cell type specific marker genes. Single gene deconvolution was performed by identifying individual marker genes of each cell type, which were used to estimate the relative proportion of each cell type with respect to the remaining markers.

### Benchmarking cell population proportion estimation on the CellBench dataset

The CellBench dataset provides gene expression profiles obtained from sequencing titrated RNA mixtures from three human lung adenocarcinoma cell lines (H1975, H2228, HCC827), as well as single cell RNA profiles from each cell line. Sequencing was performed using either plate based (CEL-Seq2 or Drop-Seq) or droplet based (10x Chromium and Drop-seq Dolomite) protocols. The proportion of RNA from each cell line was recorded for each mixture and defines a baseline for methods aiming to computationally estimate the RNA percentages. We trained scProjection using the RNA mixtures as inputs *b*_*n*_ and the single cell data as the atlas *S*. We treated the scProjection estimates 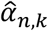 as our predictions of abundances for each cell type. We then compared scProjection-based deconvolution against other methods as described above (Supplementary Figure 8).

### Benchmarking cell population proportion estimation on the ROSMAP-IHC dataset

To provide a more challenging and realistic deconvolution benchmark, we used the ROSMAP-IHC dataset consisting of 70 bulk RNA samples of the dorsolateral prefrontal cortex (DLPFC), an scRNA-seq atlas derived from the DLPFC, and cell population proportions estimated using IHC from adjacent samples to those samples used for sequencing. The bulk RNA, reference single cell atlas and cell population proportions were collected and estimated in three different studies, thus introducing technical and biological variability between data modalities that does not exist in the CellBench study. We trained scProjection using the RNA mixtures as inputs *b*_*n*_ and the single cell data as the atlas *S*. We treated the scProjection estimates 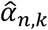 as our predictions of abundances for each cell type. We then compared scProjection-based deconvolution against other methods as described above (Supplementary Figure 9, 10). Furthermore, for each proportion estimated by scProjection we assign a confidence score indicating the certainty of the mixture being assigned to a specific cell type (Supplementary Figure 11).

### Prediction of cell population using scProjection

From scProjection’s estimates of cell population specific abundances, treated as probabilistic class assignments, the class with maximal probability is assigned as the cell population label for each sample.

### Cell annotation with *k*-NN label transfer

After estimating the projection of a mixture onto a single cell atlas the projected mixture is labeled based on annotations in the single cell atlas of its 5-nearest scRNA-seq neighbors.

### Zonated gene expression scoring

For each gene we compute the distance from an idealized zone-specific measurement as the difference between *gene*_*ideal*_= (1,0,0,0,0) and the computed gene zonation score vector. A threshold was set based on the 75^th^ quantile of the resulting scores to compare the number of zonated genes across methods.

### Constructing a gold standard set of zonated goblet expression patterns based on clumps

Manco et al.^20^ sequenced ‘clumps’ consisting of multiple physically-proximal cells from partially-dissociated intestinal villi. Using scProjection, we performed expression deconvolution and identified enterocyte-goblet clumps that contained both enterocytes and goblet cells, using a single cell atlas of enterocytes^19^ and goblet cells^20^. Based on our zonated enterocyte expression patterns (**Fig. 4b**), we predicted the zone of each enterocyte-goblet clump based on the projection of the enterocyte-goblet clump onto the enterocyte single cell atlas. Because the goblets in the enterocyte-goblet clumps are physically proximal to the enterocytes, we then assumed the goblets in each enterocyte-goblet clump was from the same zone as the projected enterocyte. For each zone, we identify all enterocyte-goblet clumps from that zone, project the enterocyte-goblets to the goblet single cell atlas, and average across all such projections to estimate zone-specific goblet expression.

