## Supplementary figures for "Projecting clumped transcriptomes onto single cell atlases to achieve single cell resolution"

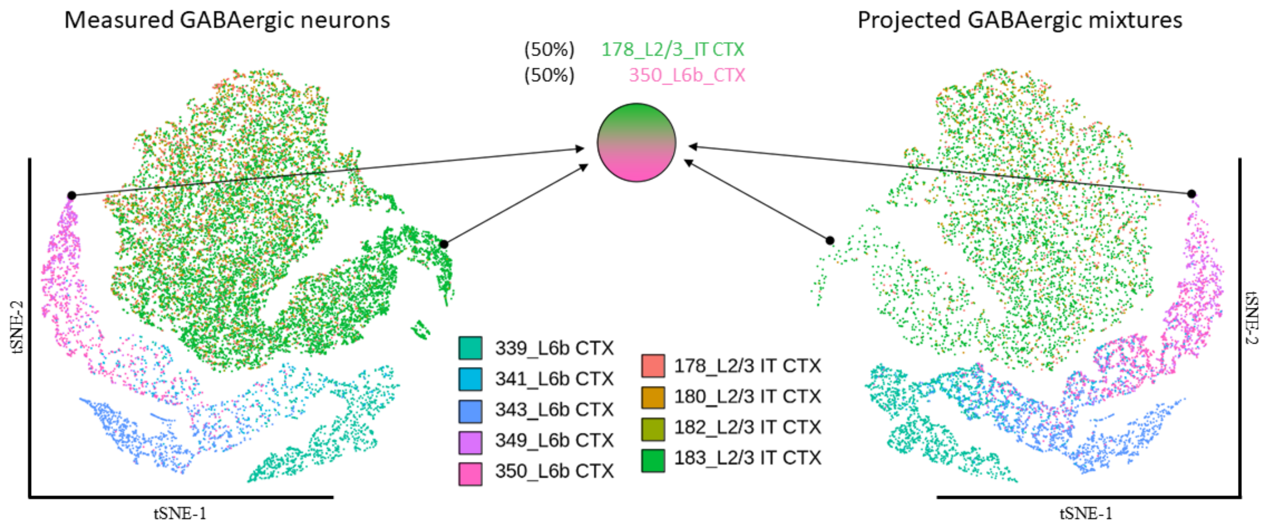

**Fig S1: Simulated mixed RNA samples from L2/3 and L6b neuronal populations.** Simulated samples were generated by selecting pairs of cells, one from each population, and adding them together. Samples were projected back onto the two populations to determine how close projections were to the originally selected cells. Shown is an example of a single mixture generated by selecting two cells (from the tSNE plot on the left), and projecting them back onto those same populations (to the tSNE plot on the right). tSNE plot visualizes the measured GABAergic gene expression patterns (left) and projected mixtures (right) in the same low dimensional space (with a reversed tSNE axis on the right for visualization symmetry).

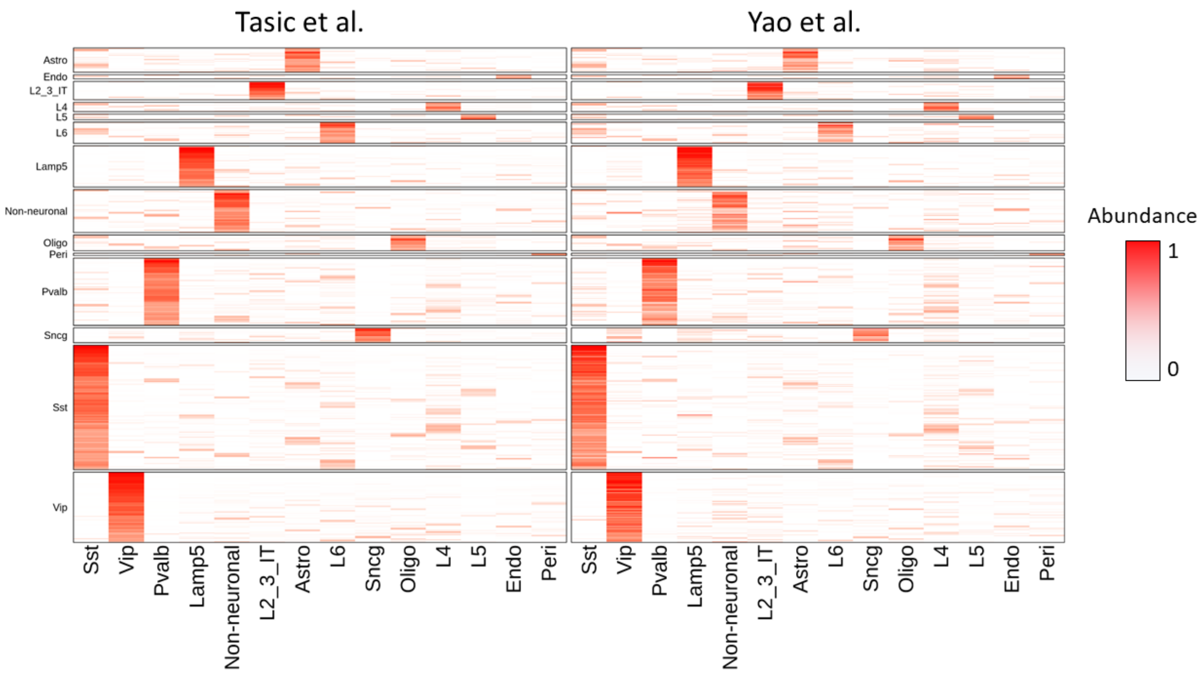

**Fig S2: Estimated cell type abundances using multiple atlases of the mouse cortex.** Heatmaps visualize the abundance of each cell type (columns) for each PatchSeq sample (rows) based on training scProjection using either the Tasic et al. atlas or a recent atlas from Yao et al.

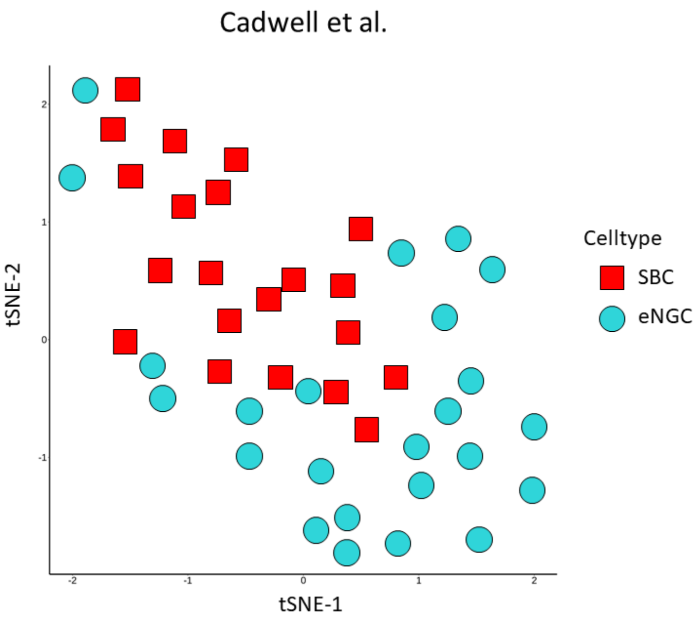

**Fig S3: Embeddings of SBC and eNGC neurons.** tSNE plot visualizes the separation of SBC and eNGC neurons from Cadwell et al. projection by scProjection.

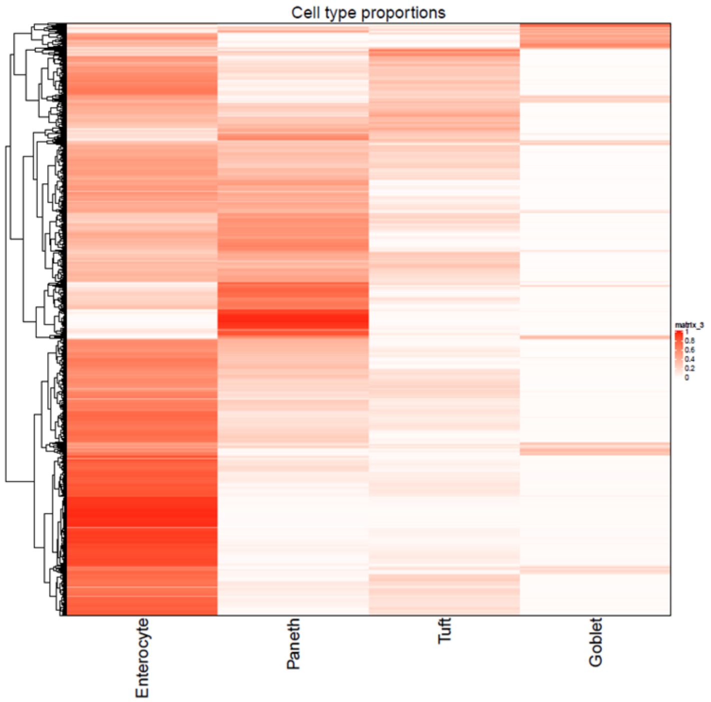

**Fig S4: Estimated cell type abundances for clump-seq samples.** Heatmap visualizes the cell type abundances of each cell type (columns) for each clump (rows) in Manco et al.

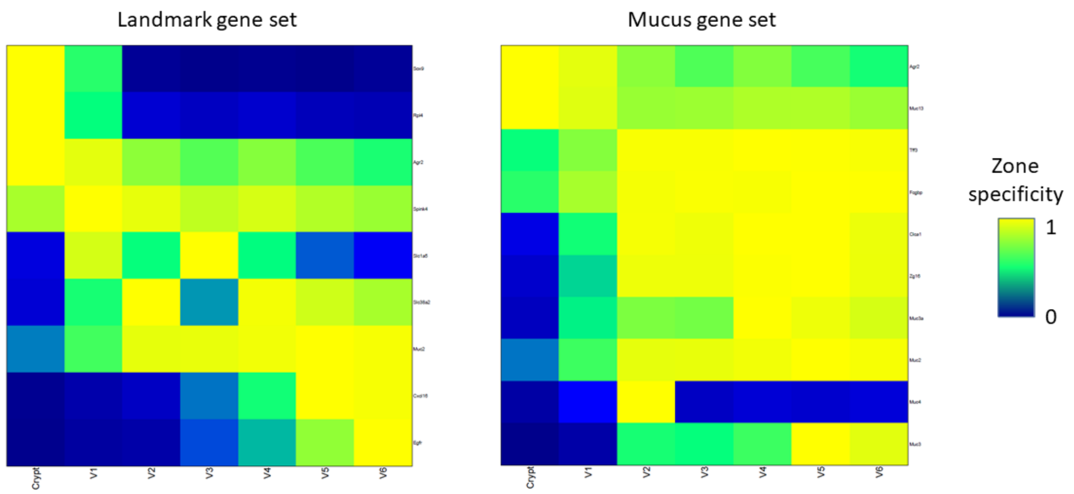

**Fig S5: Zonated expression of projected goblet clumps.** Heatmaps visualize the zonation pattern for goblet specific expression of landmark genes (left) and mucus genes (right), where yellow indicates increased zone specific expression.

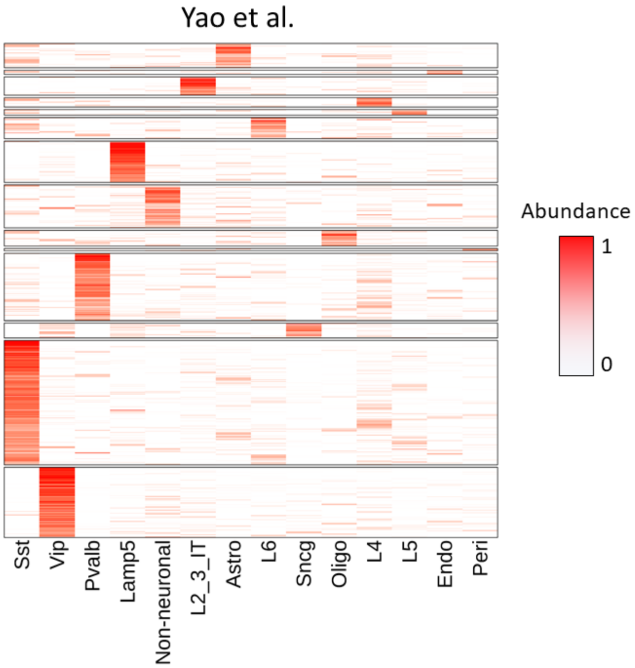

**Fig S6: Estimated cell type abundances for mouse PatchSeq data.** Heatmap visualizes the cell type abundances of each cell type (columns) for each neuron (rows) in the Allen mouse PatchSeq data.

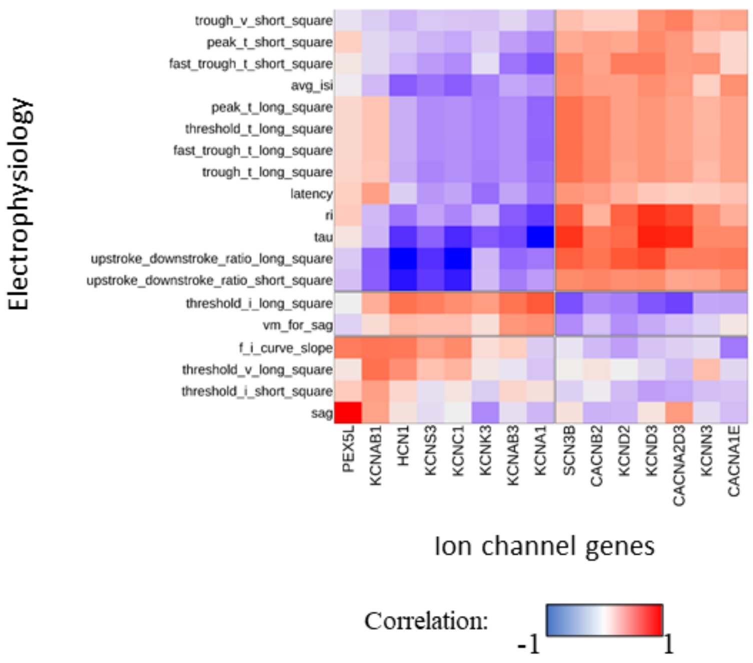

**Fig S7: Correlation of ion channel genes with electrophysiology features.** Heatmap visualizes the correlation of the most variable ion channel genes that play a role in neuronal signaling (columns) with electrophysiology features (rows).

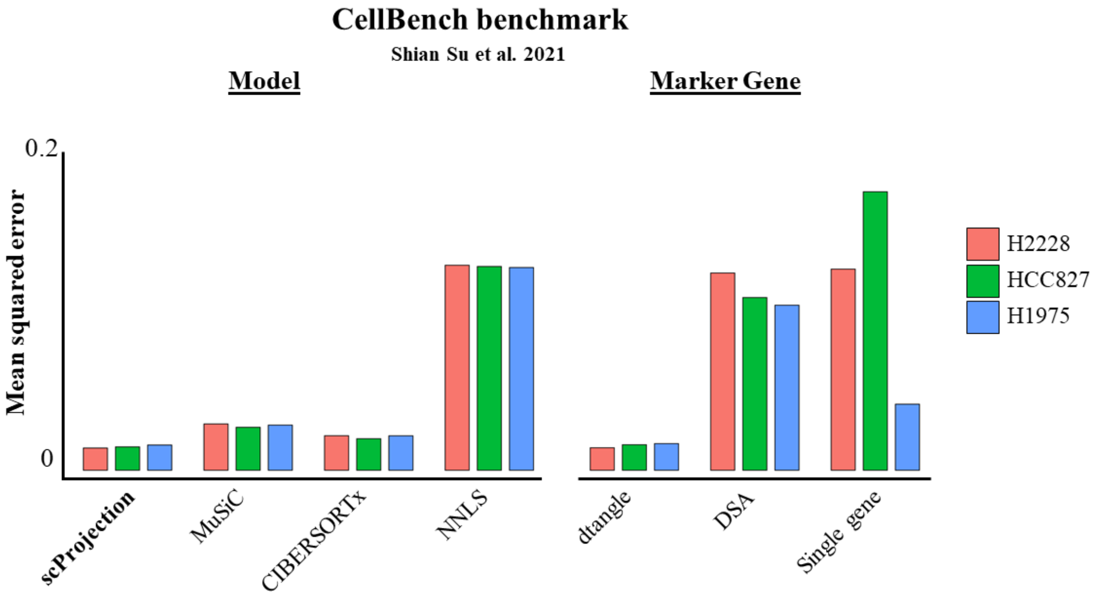

**Fig S8: Benchmarking of deconvolution methods on CellBench.** Barplots indicate the error in predicted cell type abundances compared against the ground truth for each deconvolution method.

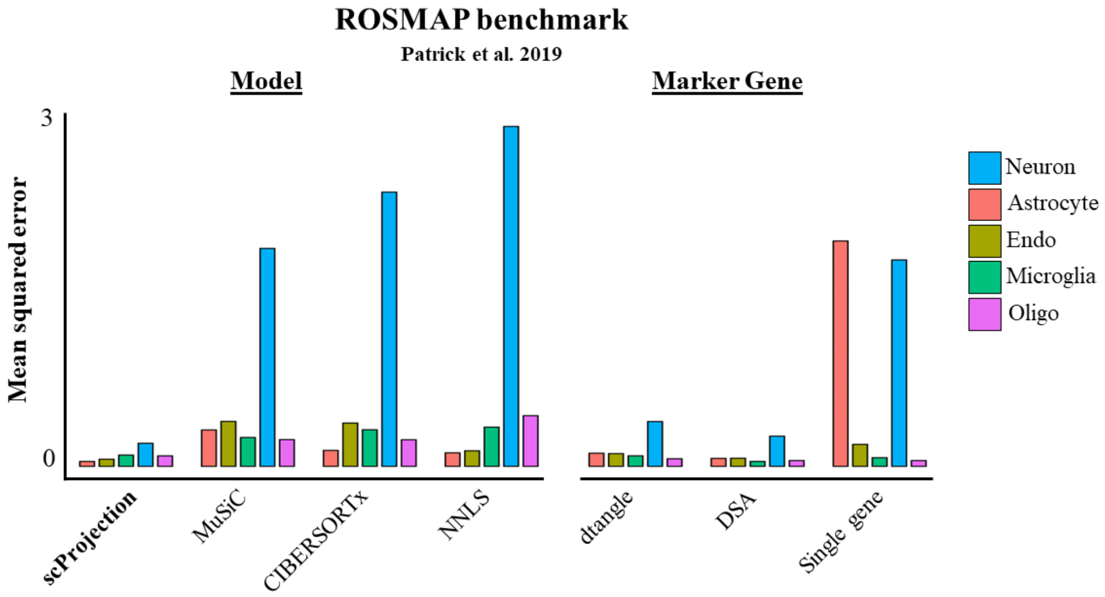

**Fig S9: Benchmarking of deconvolution methods on ROSMAP.** Barplots indicate the error in predicted cell type abundances compared against the ground truth for each deconvolution method.

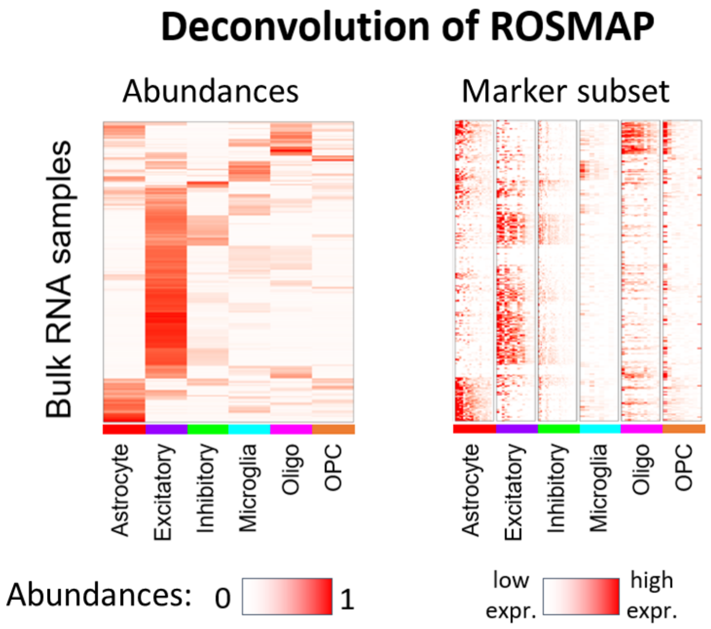

**Fig S10: Cell type abundances and marker gene expression of ROSMAP bulk samples.** Left heatmap indicates the estimated abundances of cell type (columns) for each bulk sample (rows), and the right heatmap shows the expression of the top marker gene expression (columns) for each cell type.

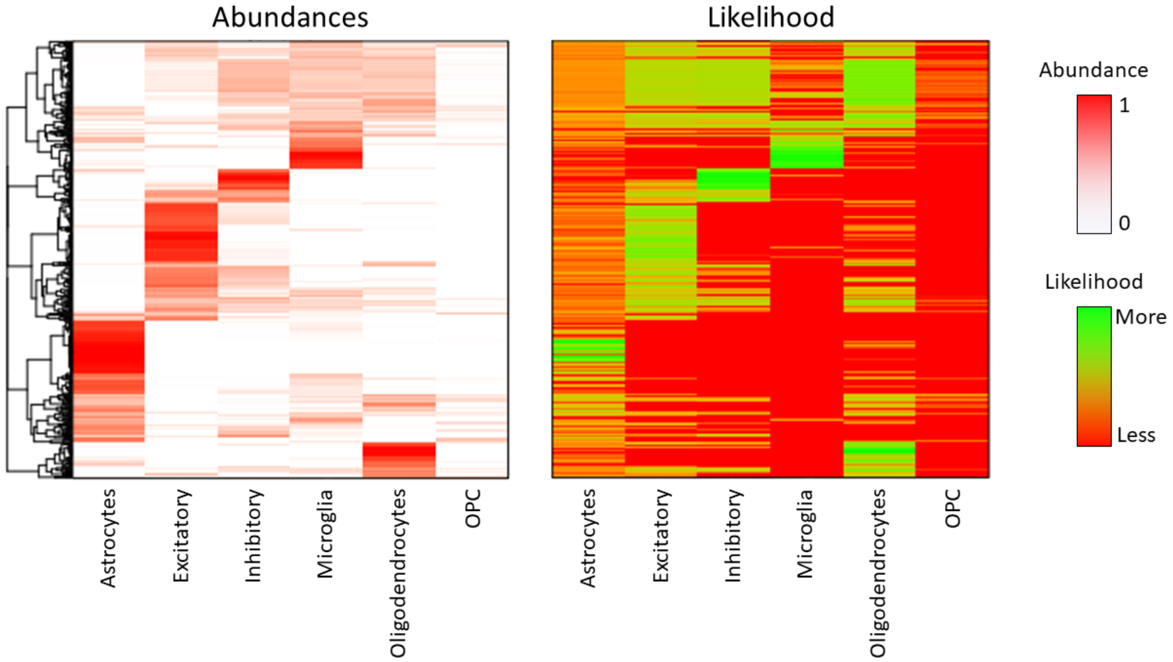

**Fig S11: Estimated cell type abundances and likelihood for ROSMAP bulk samples.** Left heatmap indicate the estimated abundances of each cell type (columns) for each bulk sample (rows), and the right heatmap shows the likelihood of each abundance value for each cell type (columns).

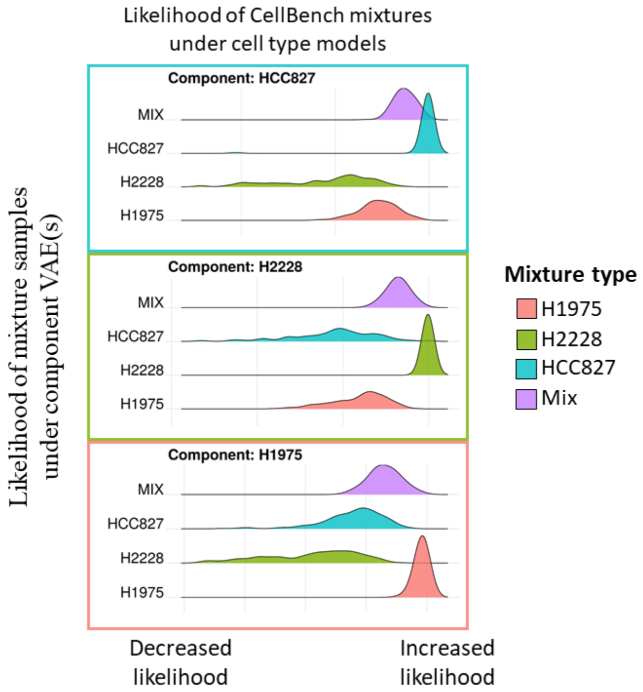

**Fig S12: Likelihood of CellBench mixtures under each component VAE in scProjection.** Density plots for each component VAE (bounded boxes) indicate the likelihood of each mixture type under the corresponding cell type model trained by scProjection.

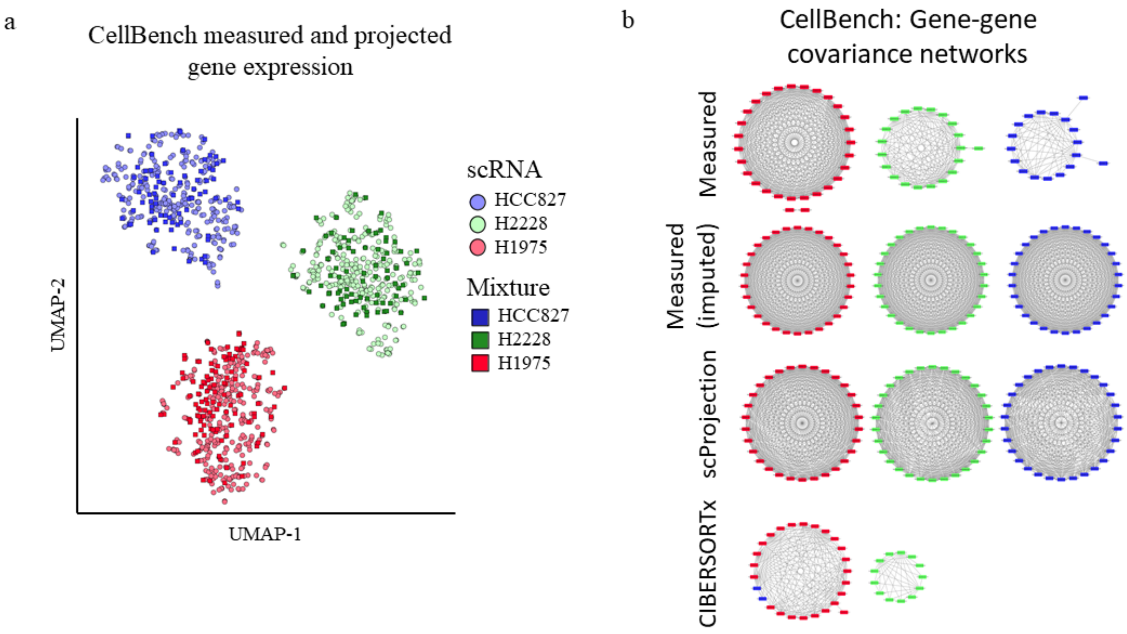

**Fig S13: Validation of projected CellBench mixtures.** (**a**) Left tSNE plot visualies the measured scRNA-seq data overlayed with the projected mixtures for each cell type. (**b**) Gene-gene correlation networks indicates the differences in conservation of gene structure by scProjection and CIBERSORTx.

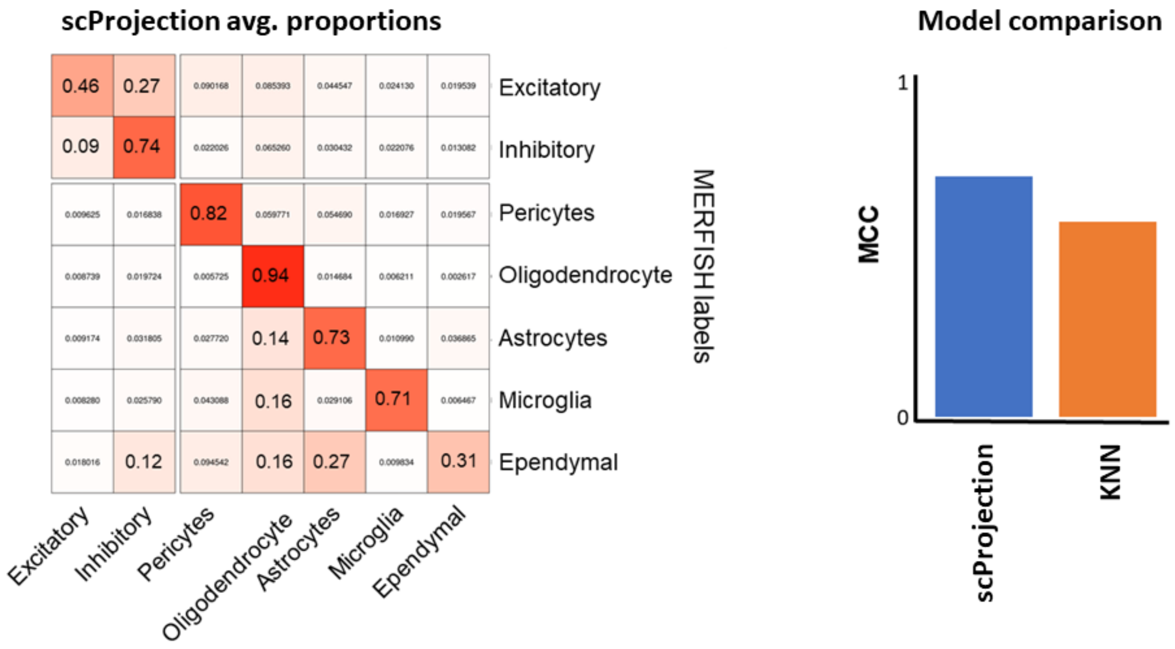

**Fig S14: Validation of scProjection cell type annotation on MERFISH.** Heatmap visualizes the average abundance for each cell type called by scProjection on the columns and the label annotated in the original MERFISH study. The barplot shows the classification performance of scProjection against a k-nearest neighbors approach which labeled based on neighborhood proximity of the MERFISH data mapped onto the scRNA-seq atlas.
