## Supplementary notes for "Projecting clumped transcriptomes onto single cell atlases to achieve single cell resolution"

### Supplemental Note 1

As a first step, we performed experiments to determine the extent to which scProjection can identify the primary cell type of an RNA sample. Specifically, we benchmarked the deconvolution performance of scProjection utilizing recent bulk RNA-seq studies for which the proportions of each cell type in the mixed RNA samples were experimentally determined[16,17]. Our first benchmark is CellBench[16], a dataset where mixed RNA samples were experimentally constructed by mixing RNA from three human lung adenocarcinoma cell lines and varying either the relative concentrations of RNA content or the numbers of cells. With an ideally matched scRNA-seq atlas, the deconvolution of CellBench mixtures was not a challenging task and most methods estimated the proportions near perfectly (avg. acc: 0.95) (**Supplementary Fig. 8**). The second benchmark is the ROSMAP dataset[17], consisting of 70 bulk RNA and 80,660 scRNA-seq samples from the dorsolateral prefrontal cortex, where abundances per cell type were estimated based on immunohistochemistry (IHC). The ROSMAP dataset presented a more challenging task than CellBench due to increased technical and biological variation between the bulk RNA samples and the reference single cell atlas. For ROSMAP, scProjection clearly performs best across all tested methods (MSE: 0.04 compared to median MSE: 1.3 for other methods) with respect to estimating cell type proportions of each bulk sample. In contrast, the remaining model based methods ( methods which do not need marker gene sets per cell type) overestimated the neuronal content of each sample and underestimated the rarer non-neuronal cell types (**Supplementary Fig. 9, 10**). A novel aspect of scProjection is the ability to compute, per sample, an estimate of the likelihood for each cell type proportion that enables the identification of samples with low concordance to the atlas (**Supplementary Fig. 11**) and higher error in reconstruction of the original mixture measurement.

**Supplemental Note 2:**

We first used the CellBench[44] benchmark to validate that projections of mixed RNA samples to individual cell populations yield cell states that resemble the single cells used to train scProjection. CellBench is a dataset which consists of scRNA-seq datasets generated on three human lung adenocarcinoma cell lines (H1975, H2228, HCC827), as well as bulk RNA mixtures of all three cell lines combined at varying magnitudes. We used scProjection to project 636 mixed RNA samples to each of the composite H1975, H2228, and HCC827 cell populations. scProjection estimated gene expression profiles and likelihood (**Supplementary Fig. 12**) for each of three cell lines per input for all 636 RNA mixtures. Projections were highly correlated (rho>=0.98) with the average measured scRNA-seq profiles for each cell line, suggesting projections globally look similar to the single cell data. We also demonstrated that our projections retain the gene co-expression networks exhibited in the measured single cell atlas after imputing for gene expression dropout (**Supplementary Fig. 13**) as compared to the deconvolution method CIBERSORTx[34]. We saw similar results on another benchmark, the ROSMAP-IHC dataset[17], where projections of 70 bulk RNA samples onto five cell types were also similar to the cell type averages (rho = 0.91)despite the higher degree of technical and biological variation. scProjection therefore projects cell population-specific expression profiles that are consistent with the single cell measured profiles from the reference single cell atlas.

**Supplemental Note 3**:

Having validated scProjection's predictions of the dominant cell type from a single sample, we next assessed scProjection for the ability to impute genome-wide expression measurements using a limited set of marker genes. In a study of neurons from the hypothalamic preoptic region of the mouse brain, Moffit et al. assayed 155 marker genes across millions of neurons using MERFISH, and generated a paired scRNA-seq cell atlas[27]. Using the scRNA-seq cell atlas, for each individual cell population, we performed a series of experiments where we randomly sampled a cell from the atlas, extracted the expression levels of only the 155 marker genes used for MERFISH, and used scProjection to impute ~4000 genes from the 155 marker genes. We found the projected and measured gene expression patterns correlated well and the predicted cell state agreed with the original scRNA-seq measurement (Spearman rho=0.63, p=2e-8)**.** We also found that scProjection could identify the correct cell type in 0.78 of cells (**Supplementary Fig. 14**). These results suggest scProjection can be used to impute genome-wide expression profiles based only on marker gene expression.
